# Computationally assisted design and selection of maneuverable biological walking machines

**DOI:** 10.1101/2020.09.01.278622

**Authors:** Jiaojiao Wang, Junehu Park, Xiaotian Zhang, Insu Park, Evin Kilicarslan, Yongdeok Kim, Rashid Bashir, Mattia Gazzola

**Author notes:** Electronic address (Rashid Bashir). Electronic address.

## Abstract

The intriguing opportunities enabled by the use of living components in biological machines have spurred the development of a variety of muscle-powered bio-hybrid robots in recent years. Among them, several generations of bio-hybrid walkers have been established as reliable platforms to study untethered locomotion. However, despite these advances, such technology is not mature yet, and major challenges remain. This study takes steps to address two of them: the lack of systematic design approaches, common to bio-hybrid robotics in general, and in the case of bio-hybrid walkers specifically, the lack of maneuverability. We then present here a dual-ring biobot, computationally designed and selected to exhibit robust forward motion and rotational steering. This dual-ring biobot consists of two independent muscle actuators and a 4-legged scaffold asymmetric in the fore/aft direction. The integration of multiple muscles within its body architecture, combined with differential electrical stimulation, allows the robot to maneuver. The dual-ring robot design is then fabricated and experimentally tested, confirming computational predictions and turning abilities. Overall, our design approach based on modeling, simulation, and fabrication exemplified in this robot represents a route to efficiently engineer biological machines with adaptive functionalities.

## I. INTRODUCTION

Building with living cells is an exciting avenue towards the synthesis of fundamental biological principles and conventional engineering design [1–5]. In this context, biological machines have become a prominent paradigm to explore this synergy, in the pursuit of both novel applications and fundamental understanding. In such bio-hybrid systems the biological component can provide actuation, sensing and even computing abilities [6], while artificial elements provide the organizational and structural template [7, 8]. This is typically implemented through an elastic, engineered scaffold (the ‘skeleton’) around which cells grow, self-organize and coordinate their activities, resulting in higher-order functionalities as a combination of internal processes and interactions with the environment [2, 5, 9, 10].

Over the past decade, this design paradigm has led to bio-integrated soft robots (biobots) that can grip, pump, swim or walk in response to external stimuli (light, mechanic/fluidic pressure, electric fields), providing a glimpse into the potential of this technology [6, 11–22]. Among these prototypes, untethered walking biobots in the millimeter/centimeter size range, have emerged as reliable platforms to explore and test new cell manipulation and fabrication protocols, design motifs and integration strategies in a consistent setting. Several generations of walkers have been demonstrated [11, 12, 23, 24]. Their designs incorporate flexible hydrogel skeletons fabricated with stereolithographic 3D printing [25], and living contractile muscle tissue engineered *in vitro* [26]. These muscle constructs can generate millinewton contraction forces leading to walking speeds up to ~2 body-lengths per minute (~0.5mm sec^−1^) [24]. Beside locomotion, biological walkers have also enabled the investigation of other bio-hybrid functionalities such as self-healing, matrix degradation, cryopreservation, and strengthening through exercise [12, 24, 27–29].

Despite these early successes, current designs exhibits several limitations. Here we focus on two of them: firstly, bio-hybrid walkers can only perform unidirectional locomotion and are incapable of turning, rotating or altering their trajectory; secondly, although progresses have been made in the computational forward design and optimization of bio-hybrid robots [6, 24], there is still a lack of systematic design approaches. As a consequence, from a pool of candidate, intuition-originated designs, often only one or two are actually fabricated and tested due to the time consuming nature of these experiments and their low success rate. Moreover, without reliable numerical tools and metrics to *a priori* assess the performance of these potential designs, the selection criteria is ultimately somewhat arbitrary.

Here, motivated by the long term goal of achieving adaptive behavior in bio-hybrid robotic systems, we focus on maneuverability while expanding the role of computational design for rapid prototyping and selection. The result is a versatile dual-ring bio-hybrid robot capable of walking straight or turning in response to controllable external electrical stimulation.

We utilize a recently introduced simulation approach for soft, heterogeneous musculoskeletal architectures [30, 31] to model and test a variety of intuitive designs. Based on this computational analysis, a design striking a balance between walking speed, robustness and turning ability is selected. This design employs two independent actuators attached to a 4-legged skeleton, allowing for localized control and tunable behavior. Finally, the selected prototype is fabricated and characterized, verifying predicted performance. Our results pave the way to advance the design, fabrication, and optimization of more complex multi-functional bio-hybrid systems in the future.

## II. RESULTS AND DISCUSSION

### A. Intuitive designs and computationally assisted selection

Previous bio-hybrid walking robots (Figure 1b) consist of a hydrogel scaffold made of two pillars (legs) and a connecting bridge [12, 24, 27–29]. Skeletal muscle tissue is shaped so as to wrap around the pillars, an architectural motif reminiscent of the muscle-tendon-bone relationship found *in vivo*. Muscle contractions bend the legs inwards, flexing the bridge and storing elastic energy that is subsequently released during the muscle relaxation phase. As a result, cyclic frictional forces are generated at the leg-substrate interface. Symmetry is broken by having a leg shorter than the other, giving rise to net unidirectional forces, thus forward locomotion.

**FIG. 1:**
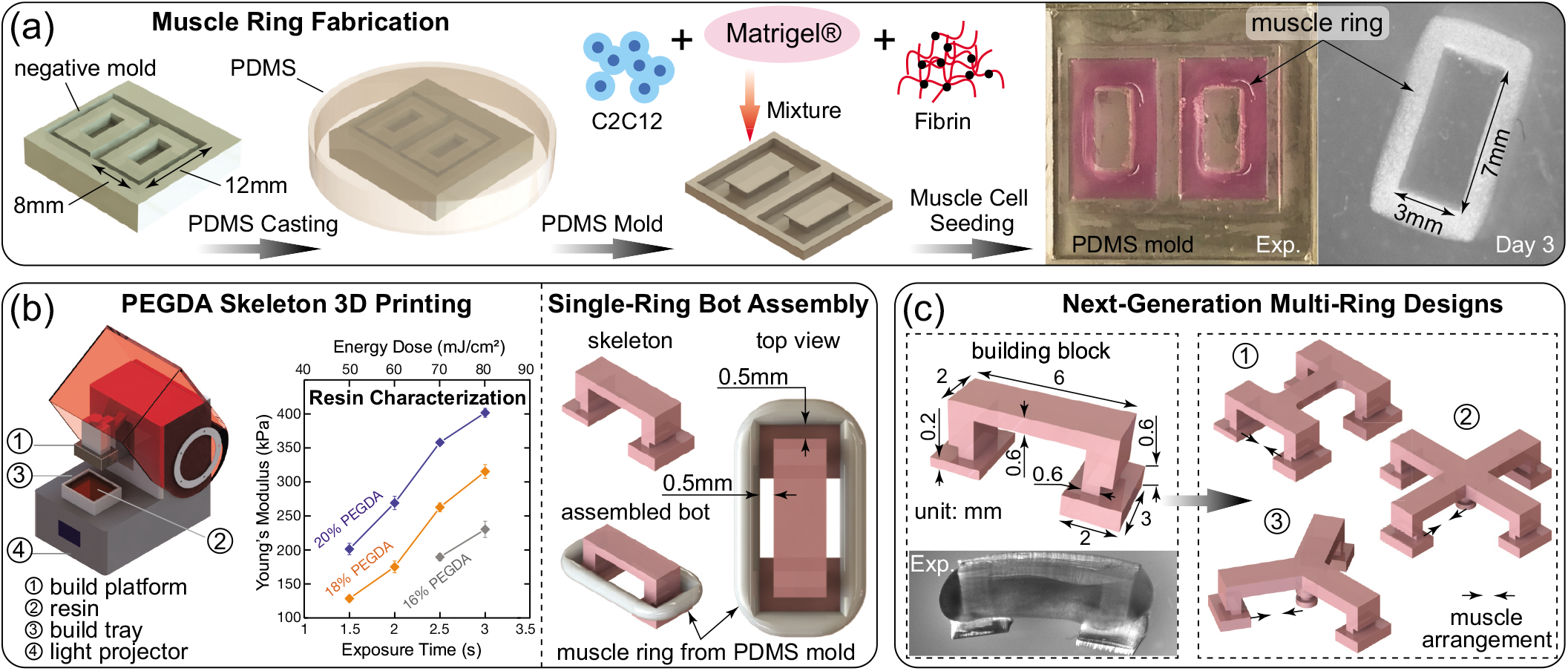
Biofabrication process of bio-hybrid walking robots. (a) C2C12 muscle cells and ECM proteins are seeded in the PDMS mold, casted from a plastic negative mold, to form 3D muscle rings. (b) The skeleton of the bio-hybrid robot is printed using a DLP 3D printer and well-characterized PEGDA resin. Compacted muscle rings from (a) are then assembled on the printed structure. (c) A single-ring biobot is fabricated to characterize the muscle ring force output. Based on the established single-ring architecture, three intuitive multi-ring designs are proposed.

Then, to achieve controllable turning maneuvers, we need to move away from single-muscle ring constructs, and instead consider multiple-muscle layouts that can generate *lateral* net forces as well. This augmented design space lends itself to a number of potential solutions. Based on our previous understanding of the single-ring bio-hybrid walker [11, 12], three next-generation bio-hybrid walker devices are proposed: dual-ring, tri-ring, and quad-ring biobots (Figure 1c). Next, to enable a rational selection process, the three intuitive designs are modeled as assemblies of Cosserat rods [30, 31] and numerically evaluated.

To inform a muscle model able to recapitulate realistic force outputs, we started by fabricating a testbed consisting of one engineered muscle ring and a two-pillars soft scaffold. Muscle rings were formed by embedding myoblasts in an ECM solution and casting into a PDMS mold for compaction (Figure 1a). The scaffold was 3D-printed using a well-characterized digital-light-projection-based printer (Figure 1b – refer to the experimental section for details). Static tension (passive force) and cyclic contractions (active force) induced by electrical stimulation were characterized by measuring deflections of the pillars. Figure 2b shows our measurements relative to 5 samples, for different stimulation frequencies. This characterization provides us with average muscle outputs and associated intrinsic variability. Because of experimental uncertainties relative to cell density, myotube width and alignment, the active force produced by different muscle rings of the same batch can vary more than 40%. Based on these data, we created a corresponding virtual muscle as in [6, 24]. For verification, we wrapped our model muscle around a computational scaffold with the same geometric and material properties of the experimental setup (refer to SI for more details). Upon actuation, simulated scaffold deflections (based the average motor outputs of Figure 2b) are found to be in good agreement with experiments (Figure 2a,c).

**FIG. 2:**
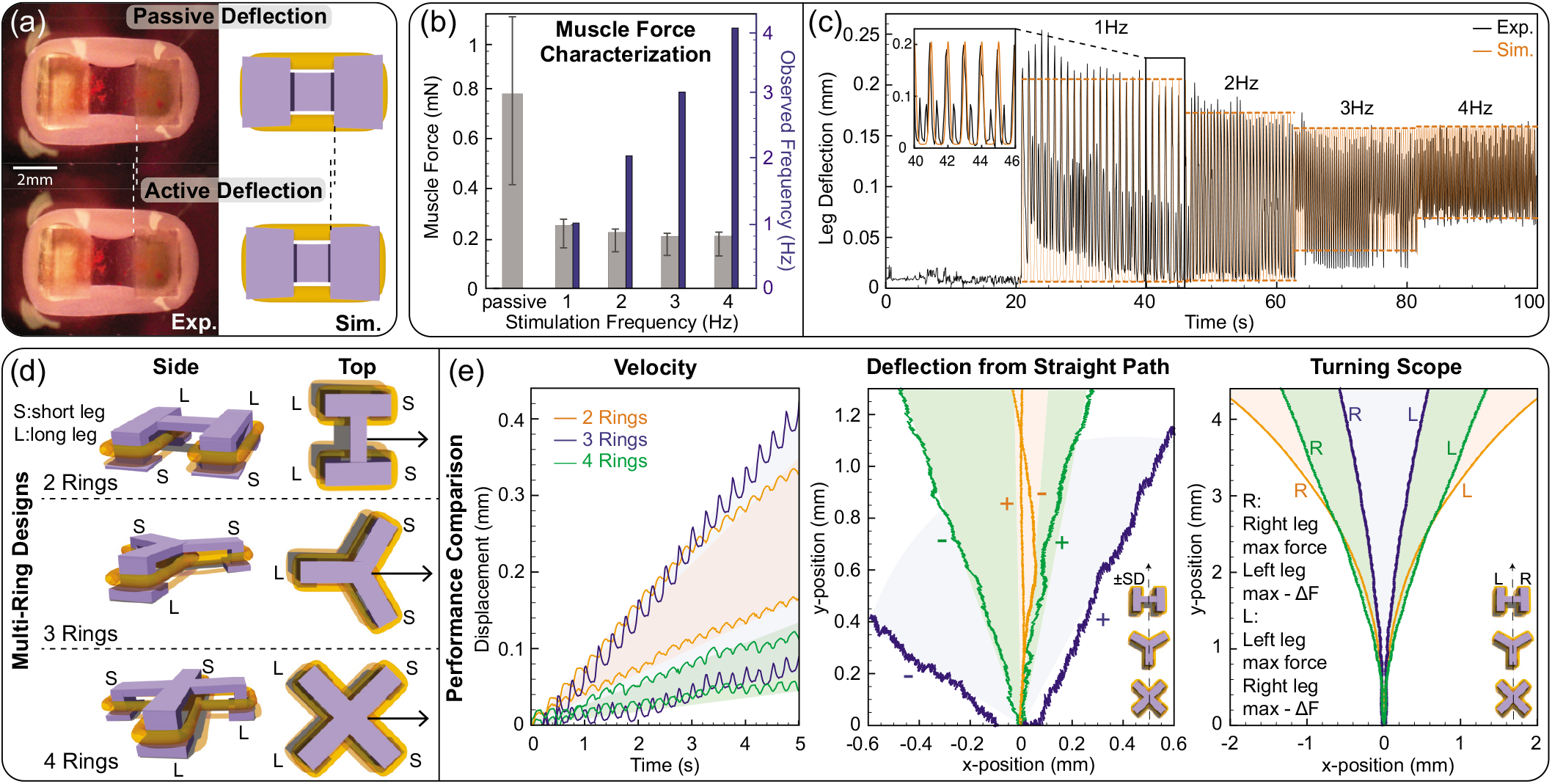
Computationally assisted selection of multi-ring biobots. (a) Illustration of passive and active deflection from experiment (left) and on the virtual reconstruction of the single-ring testbed (right). (b) Rest tension and cyclic contraction force experimentally obtained from single-ring testbeds stimulated at 1, 2, 3, and 4 Hz (mean ± SD, n=5). Observed frequencies from the experimental pillar deflection data are plotted to confirm muscle responsiveness. (c) Measured leg deflection data at different frequencies used to calibrate our muscle model, and predicted leg deflection data on the virtual testbed when average forces and observed frequencies from (b) are used as inputs. (d) Our three multi-ring biobot intuitive designs. (e) Predicted walking and turning behaviors for the three multi-ring biobots. All input forces are based on the passive force and the 4 Hz active force from (b). *Velocity:* The velocity range for each design is obtained through 7 simulation samples with a passive force variation (mean ±10% SD, 752 ± 33.6 *μ*N) and an active force variation (mean ±1 SD, 182 ± 49 *μ*N). The lower/upper bound for each design is produced by the following passive and active force combination: 2 Rings (+10% SD passive, +1 SD active / −10% SD passive, +1 SD active), 3 Rings: (+10% SD passive, −1 SD active / +10% SD passive, mean active), and 4 Rings (−10% SD passive, −1 SD active / −10% SD passive, mean active). *Deflection from straight path:* When uniformly electrically stimulated, and in the absence of uncertainties, the biobots walk in a straight line aligned with their initial bearing. Here we assess instead how imperfect (i.e. asymmetric due to experimental uncertainties) muscle responses affect the ability to walk straight. Biobots characterized by small deflections are less sensitive, hence more reliable in performing directional motion. Walking trajectories for each design are simulated using for all muscles the average passive force of 752 *μ*N from (b). All right legs produce active forces equal to the average value of 182 *μ*N from (b). Left legs instead exhibit an active force of 182 *μ*N ±1 SD. This analysis is not meant to be a comprehensive uncertainty quantification study, rather a rapid sensitivity estimate, in line with the prototyping character of these simulations. *Turning scope:* To access the potential steering capabilities of our designs, we consider passive forces of 1079 *μ*N (the maximum passive force from (b)), and active forces of 239 *μ*N (the maximum active force from (b)) on one side of the bot and 89 *μ*N (maximum active force – ΔF) on the other side. The differential ΔF = 150 *μ*N has been selected to be approximately consistent with the maximum active force differential from (b). Resulting trajectories are rotated to align the robots’ initial bearing with the vertical direction, to ease comparison (raw turning trajectories reported in Figure S2).

Armed with a muscle model tailored to our biofabrication protocol and desired ring dimensions, we virtually assessed the performance of our three intuitive designs (Figure 2d,e). The goal is to select the prototype that best compromises between locomotion speed, turning abilities and performance robustness. Indeed, as underscored in Figure 2b, the behavior of our biological actuators can significantly vary across samples, thus a good design must be able to perform reliably in the face of uncertainty. The three designs were then tested assuming 4 Hz stimulation frequency, for different combinations of passive/active forces to account for variable, asymmetric motor outputs. As can be seen in Figure 2e, the 3-ring biobot is the fastest (although the one characterized by the largest spread), closely followed by the 2-ring model. Nonetheless, the 2-ring biobot outperforms all other designs when it comes to straight walking or turning, exhibiting the highest level of reliability and sharpest turns. Thus, based on this preliminary analysis, the dual-ring biobot was selected for fabrication and experimental validation.

### B. Symmetric stimulation and forward walking

As the design of the biobot’s layout is crucial to its performance, so is the design of the actuation strategy. Then, after settling on the dual-ring prototype, we proceed with the design and computational assessment of the electric field (E-field) stimulation setup. First, we focus on forward straight walking and consider a system made of two parallel, longitudinal platinum electrodes of 20 mm length, set 20 mm apart, and with applied voltage difference of 20 V. The electrodes are immersed in physiological solution within a petri dish of 35 mm radius (Figure S4a). When unperturbed, this setup generates an electric field symmetric about the horizontal midplane, and approximately uniform at the center of the domain. Then, a virtual dual-ring biobot is placed within the petri dish at three different locations: at the center of the stimulation setup and aligned with the horizontal symmetry axis, as well as at distances of 1 mm and 2 mm from the symmetry axis in the y-direction (Figure 3e). We seek to ascertain how the biobot presence and location affect the local electric field ‘felt’ at the two muscle rings, and whether symmetric stimulation is approximately preserved for forward walking. COMSOL^®^ simulations (Table S2, Figure S3) reveal that symmetric stimulation is indeed preserved (Figure 3e), and that this setup is suitable to test and control straight walking (Figure S5).

**FIG. 3:**
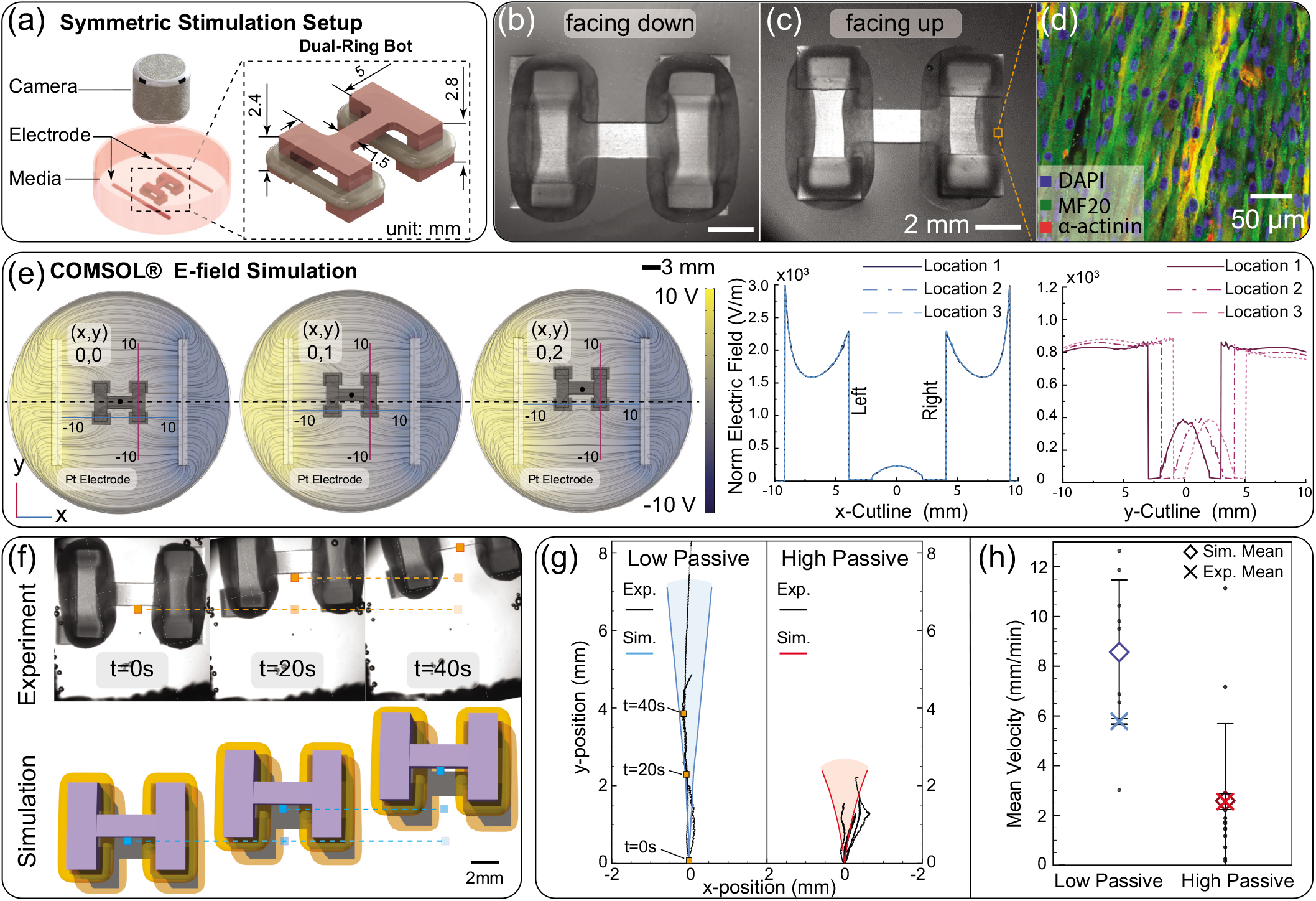
Straight walking and symmetric electrical stimulation. (a) Illustration of the symmetric electrical stimulation setup with two 20 mm Pt electrodes. The dual-ring biobot dimensions are displayed on the CAD image. (b) Bright-field image of the dual-ring biobot facing down. (c) Bright-field image of the dual-ring biobot facing up. (d) Confocal imaging of the muscle tissue expressing MF 20 myosin heavy chain and *α*-actinin (right) shows myotube distribution and muscle striation. (e) The symmetric stimulation setup is modeled in COMSOL^®^. The electric field strength distribution along x-axis and y-axis for three different bot locations are simulated and compared. (f) Walking of the dual-ring biobot is illustrated by three still images from a representative walking video and compared with computational simulated results (passive force 150 *μ*N, measured from the dual-ring biobot in this panel; active force 182 *μ*N, the average value at 4 Hz from Figure 2b). (g) Experimental walking trajectories of six different dual-ring biobots are sorted into two groups, the high passive force group and the low passive force group, and compared with simulated results. In simulation, we account for muscle variations by including the largest active force difference from the Figure 2b dataset, with one muscle ring characterized by the maximum active force of 239 *μ*N and the other by the minimum active force of 114 *μ*N. The passive forces used are the maximum values of experimentally measured for each group (150 *μ*N for the low passive force regime and 1100 *μ*N for the high passive force regime). (h) Walking velocities of the actual biobots fabricated and tested are sorted in the high /low passive force groups and compared to model predictions. The virtual dual-ring biobots are simulated using the experimentally observed passive forces in each group (min, average, max: 0, 75, 150 *μ*N for the low passive force group and 550, 825, 1100 *μ*N for the high passive force group) and minimum, average, and maximum active forces (114, 182, 239 *μ*N) at 4 Hz from the single muscle ring characterization data of Figure 2b. Average simulation and experimental velocities ± SD are indicated in the plot for comparison.

With a better understanding of the E-field distribution, we proceeded to fabricate the system and deploy our dual-ring biobot to assess its functionality (Figure 3a,b). The dual-ring structure was fabricated with the same protocol as the single-ring biobot: stereolithographic 3D printing and tissue engineering based on myblasts and ECM solution. In order to verify the myogenic health of the tissue-engineered muscle rings, immunohistochemical staining was performed (Figure 3d). Differentiated myotubes were found to be distributed in the tissue and aligned along the muscle ring. The observed muscle striations indicate the tissue’s capability of generating force for locomotion (Figure 3d)[32]. Before testing their walking performance, we further characterized muscle force outputs of every sample, by turning the robot upside-down to measure cyclic leg deflections for different stimulation frequencies (Figure 3c). These data were then used to aid the comparison with simulations, and in particular to sort biobots based on muscle passive forces, as discussed in the following.

Once placed in the petri dish and stimulated at 4 Hz frequency, the biobot dynamics finally resulted in forward directional walking (Figure 3f). The fore/aft geometrical asymmetry of the dual-ring biobot induced directed locomotion towards the side of the longer pillars with a mean average velocity ranging from 2.3 mm min^−1^ to 6 mm min^−1^, which is approximately one body length per minute (Figure 3h). Instantaneous velocities reveal an initial acceleration phase of approximately 10 s, followed by a plateau (Figure S6a). We attribute this behavior to the shifting E-field strength distribution away from the setup horizontal symmetry axis (Figure 3e). Indeed, due to intrinsic muscles variability, biobots do not walk perfectly straight, but present slight turning angles, thus moving in both x- and y-direction, and off the axis of symmetry. Figure 3g depicts the walking trajectories of our six functional bots, which are found to be quantitatively compatible with simulated ranges (shaded regions in Figure 3g), based on the muscle output data recorded before each experiment by turning the biobot upside-down.

Interestingly, our simulations predict an important role of muscle rings’ passive forces in dictating the walker behavior. During the maturation process, muscle rings develop an internal tension (passive force) [33] which can vary significantly across samples (Figure 2b). Then, once the muscle is applied to the biobot’s skeleton, its passive force causes the beam connecting the legs to flex. A large passive force produces a persistent, pronounced bending, which affects the legs’ contact angle and friction forces distributions, as if the biobot was ‘tiptoeing’ on the petri dish substrate. In our simulations, these excessive deformations are found to be detrimental both for speed and for reliably maintaining forward bearing. To confirm our predictions, we sorted the six fabricated robots into two groups: biobots exhibiting high passive forces (550 – 1100*μ*N) and biobots characterized by low passive forces (0 – 150*μ*N), where the passive force for each dual-ring biobot is the average of the two muscle rings. Biobots with high passive forces were found to walk at the average speed of 2.5 mm min^−1^, whereas biobots with low passive forces turned out to be more than twice as fast with an average velocity of 5.9 mm min^−1^ (Figure 3h). In addition, trajectories were confirmed to be negatively affected by high passive forces, presenting a much less consistent behavior, exhibiting sudden orientation changes and overall struggling with maintaining forward directionality (Figure 3g). This characterization underscores the utility of simulations not only to design, but also to subsequently analyze and understand the mechanisms at play.

### C. Asymmetric stimulation and rotational steering

As illustrated in the previous section, our dual-ring biobots achieve forward walking by combining symmetric actuation with asymmetric fore/aft body geometry. Then, rotational steering can be obtained by breaking left/right actuation symmetry to produce net *lateral* frictional forces, a route made possible by the presence of multiple muscles. To control this process, we design a second local stimulation strategy and setup. We consider two parallel platinum electrodes, aligned in the y-direction at a separation distance of 20 mm, and with an applied voltage difference of 20 V (Figure 4a and Figure S4b). The electrodes are 3 mm long, much shorter than in the previous setup. This generates a dipole-like electric field with strong horizontal gradients, suitable to achieve differential stimulation between the two muscles. Again, we modeled this setup in COMSOL^®^ (Table S2, Figure S3) to assess the local field strength at the legs of a virtual biobot placed at three different, representative locations/orientations (Figure 4c). Numerical results indicate E-field differences of ~2 V cm^−1^ (Figure 4c), that approximately correspond to a 50% right/left stimulation imbalance, a difference expected to suffice for steering.

**FIG. 4:**
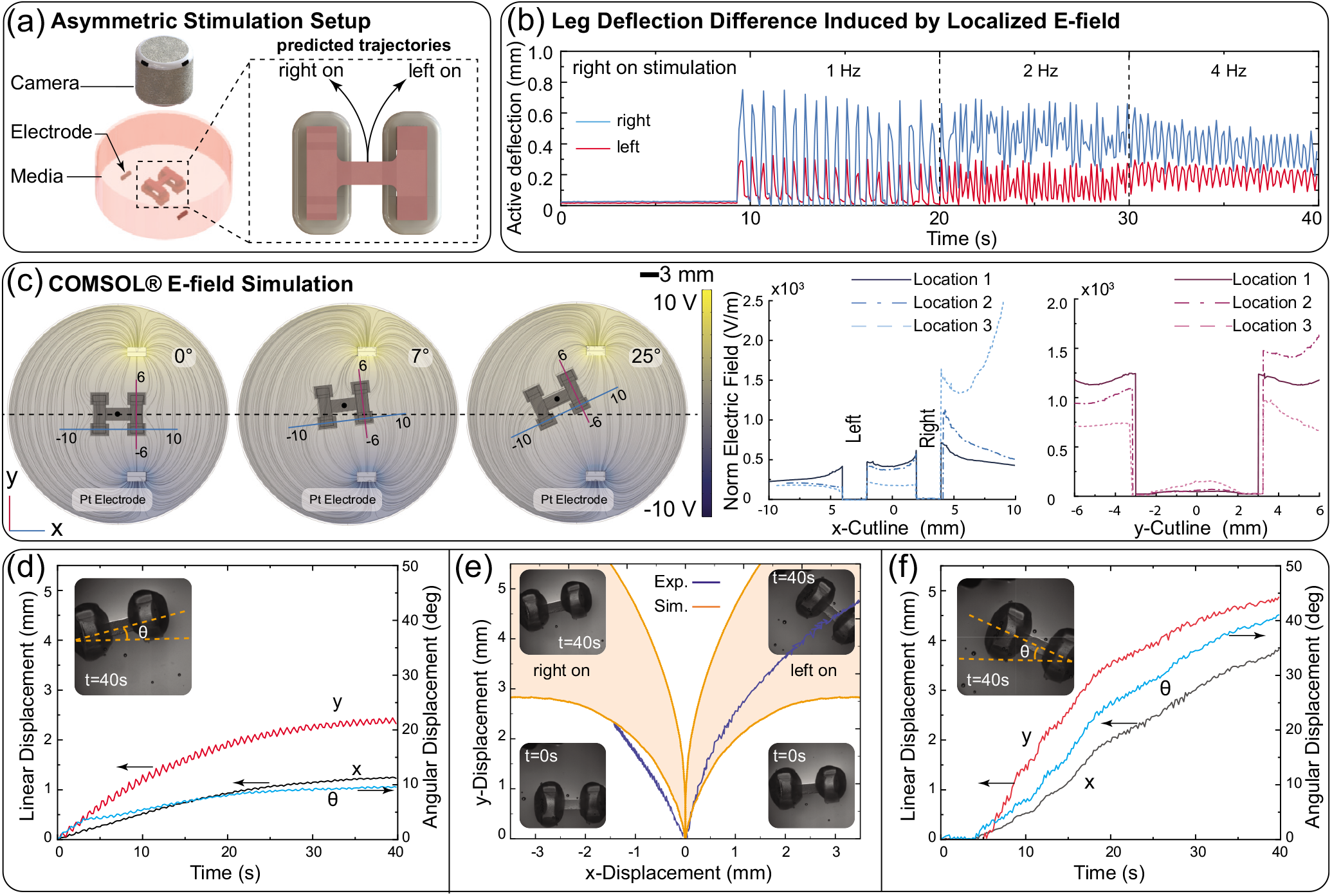
Controllable turning and asymmetric electrical stimulation. (a) Illustration of the asymmetric electrical stimulation setup with two 3 mm Pt electrodes. The expected turning directions of the dual-ring biobots are displayed on the CAD image. (b) Measured leg deflection data of left/right sides under asymmetric stimulation, which are used as inputs for simulations. (c) The asymmetric stimulation setup is modeled in COMSOL^®^. The electric field strength distribution along x-axis and y-axis for three different biobot locations/orientations are simulated and compared. (d) Linear and angular displacement versus time of the dual-ring biobot when tuning left. (e) Turning trajectories of the dual-ring biobot compared to the model predictions. The predicted ranges of turning trajectories are determined by considering two passive forces: the average passive force (752 *μ*N-lower bound) and the maximum passive force (1079 *μ*N-upper bound) from Figure 2b. The active forces are instead directly based on the measurements of panel (b) of this figure. (f) Linear and angular displacement versus time of the dual-ring biobot when tuning right.

We then fabricated the setup for testing. As in the previous section, before deploying our dual-ring biobot, we characterized the effect of differential stimulation on the muscles outputs. To this end the biobot was placed with the pillars facing up, and one leg pair aligned with the electrodes. By measuring the cyclic leg deflections induced by stimulation at different frequencies (Figure 4b), we quantified muscles’ contraction forces and their asymmetry. These data were then used as input to a biobot virtual replica, to compare simulated and experimental trajectories.

After this characterization, the biobot’s turning functionalities were finally tested. As can be seen in Figure 4e, the biobot exhibits left steering when the electrodes are initially aligned with the right leg pair. This is intuitively explained by the fact that the right side of the biobot is more strongly stimulated, producing larger contraction forces and deflections relative to the left side. As a result the right side walks faster, causing the biobot to steer to the left. As expected, a mirrored behavior and right turning is observed (Figure 4e) when the left side of the biobot is initially aligned with the electrodes. To quantify steering, the x-y position of the biobot was tracked together with its bearing and instantaneous angular velocity (Figure 4d,f). The turning speed is found to be as high as 60 degree min^−1^ during the initial phases. As the turning angle increases, the biobot tilts relative to the axis connecting the two electrodes. Consequently, the difference in E-field strength at the left/right actuators decreases, leading to the turning rate reduction reported in Figure 4d,f and Figure S6b.

Finally, we input the muscle characterization data gathered before deploying the biobot into our simulation to compare resulting trajectories. As it can be seen in Figure 4e, our numerical results are consistent with experimental observations, confirming the prediction abilities of our computational design approach. Different from the straight walking case, in the turning scenario, both experiments and simulations point to a positive role of high passive forces. Indeed, they are found to contribute to faster, consistent turning, rather than making the biobot slower and more erratic as in Figure 3g,h. A potential explanation is again related to the previously observed ‘tiptoeing’ behavior that makes the bot more susceptible to turning. If the stimulation is symmetric, random factors cause uncontrolled directional changes, with detrimental effects (Figure 3). In the turning scenario instead, the asymmetric stimulation produces a consistent right/left imbalance that dominates other random variables, making the biobot both more controllable and easier to steer. Because of the observed significant impact of passive forces, better control of muscle internal tension at the fabrication stage is then identified as an important area for further improvement.

## III. CONCLUSION

In this work we introduce a novel dual-ring biohybrid walker consisting of two independent muscle ring actuators and a 4-legged hydrogel skeleton, and demonstrate its directional walking and rotational steering abilities. This dual-ring biobot design is selected based on several performance criteria, such as walking speed, turning ability and robustness, from a set of candidate, intuitive designs modeled and evaluated numerically. After fabrication and testing, experiments are found to confirm the prediction capabilities of our simulations, thus their utility to prototype, assess and optimize prior to fabrication. This systematic approach promises to accelerate the advancement of bio-hybrid technology, by enabling the rapid development of more complex designs and functionalities, towards the long-term goal of autonomous behavior at the individual or group level [34]. The maneuvering skills herby demonstrated represent a first step in this direction. Because of its reliability and architectural features, the proposed walking platform naturally lends itself to further explore the on-board integration of miniaturized electronics as well as neuronal components [6, 35], and push the sensory-motor frontier [36].

## IV. EXPERIMENTAL SECTION

### A. Assembly of dual-ring biobots

#### 1. Fabrication and preparation of PDMS molds

In order to fabricate the PDMS molds, we first designed a negative mold using a CAD software (SolidWorks) and sent it to ProtoLabs for 3D printing. To preserve this original 3D-printed negative mold, several duplicates of this negative mold were made from Smooth-Cast^®^310 (Smooth-On). To make the PDMS molds, PDMS base and curing agent Sylgard 184^®^ (Dow Corning) were mixed with 10:1 ratio by weight and degassed in a vacuum desiccator. The mixture was then poured onto the negative mold duplicates in a petri dish and cured in 60 °C overnight. The PDMS molds were peeled off from the negative structures the next day and sterilized by autoclaving for 30 min followed by UV light sterilization in BSC for 1 h (Figure 1a).

Prior to seeding the cell-gel solution, PDMS molds were treated with 1% w v^−1^ pluronic F-127^®^ (Sigma Aldrich) dissolved in PBS. The pluronic F-127^®^ was used to reduce cell and protein adhesion to the PDMS molds. After each use, PDMS molds were sterilized in 10% bleach for 10 min, in 70% isopropanol for 10 min, and washed with DI water for 10 min. Sterilized PDMS molds can be stored in 60 °C and remain their functionalities for months.

#### 2. Cell culture of 2D myoblasts

C2C12 immortalized mouse myoblasts (ATCC) were seeded at 1E6 cells per T-75 cell culture flask. Cells were cultured in growth media (GM) consisting of Dulbecco’s modified Eagle medium (DMEM) supplemented with 10% fetal bovine serum, 1% L-glutamine, and 1% penicillin-streptomycin. Media was changed daily, and cultures were kept in incubators at 37 °C and 5% CO_2_. Once C2C12 cells reached 70-80% confluence in ~2-3 days, they were lifted using TrypLE™ Express Enzyme (1X) phenol red and centrifuged to generate a cell pellet. Exhausted cell media was aspirated out carefully, and the pellet was resuspended in a small volume of GM for counting.

#### 3. Construction of 3D skeletal muscle rings

Cultured 2D C2C12 murine myoblasts less than 10 passages were used for all the functional dual-ring biobot formation. With known cell concentration, the cell suspension was aliquoted into 15 mL conical tubes, with each tube containing 3E6 cells. The cell suspension in each conical tube was centrifuged to remove remaining media. Cell pellets were then placed on ice along with all the other reagents for forming the muscle tissue. The pellet was resuspended with 115 *μ*L GM^+^ containing 98% GM and 2% 6-aminocaproic acid, clotting buffer for fibrinogen (ACA, Sigma-Aldrich), to reach a final concentration of 1E7 cells ml^−1^. The remaining reagents were added in the following order, 6 *μ*L of 100 U ml^−1^ thrombin (Sigma-Aldrich), 90 *μ*L Matrigel (Corning), and 75 *μ*L fibrinogen (Sigma-Aldrich) at a concentration of 16 mg mL^−1^ for a total of ~300 *μ*L of cell-gel solution (Figure 1a).

To form the muscle ring, 130 *μ*L of well-mixed cell-gel solution was hastily seeded into each of the two wells of the PDMS mold due to solution coagulating quickly. Molds containing seeded muscle cells were placed in incubators at 37 °C and 5% CO_2_ for 2 h to allow for further solidification. After 2 h, wells were filled with 2 mL of GM^+^ for further cell growth and tissue formation over the course of 3 days, with media changed daily.

#### 4. 3D printing of biobot structures

The skeletal structure of the dual-ring biobot was printed using a digital light processing (DLP) 3D printer (Asiga PICO 2) (Figure 1b). The printing process functions by exposing liquid resin to a light source (light projector) to polymerize a desired layer thickness (0.2 mm). The layers stack to form the solid structure. The resin was made out of 20% v v^−1^ PEGDA 700 (Sigma-Aldrich) in DI water with 0.1% w v^−1^ photo initiator (LAP, Sigma-Aldrich) and 0.04 % w v^−1^ Sunset Yellow dye (Sigma-Aldrich) to prevent light-scattering effects. The skeletal structures of the dual-ring biobot were designed in SolidWorks and set up in the 3D printing software (Asiga Composer) to specify the layer height and exposure time, as well as the desired orientation of the build on the printing platform.

Two glass coverslips (22 mm × 22 mm) were functionalized with 3-(trimethoxysilyl) propyl methacrylate (3-TPM, Sigma-Aldrich) then taped to the building platform so the structures stayed attached during the printing process. When the printing process was finished, the structures were removed from the building platform. Prior to culturing, the printed structures were placed in a 10% bleach solution for 1 h and washed with PBS for at least 1 h to remove the dye.

#### 5. Final assembly of dual-ring biobots

Once the cell-gel mixture was compacted to a ring shape in the PDMS mold over 3 days, the muscle rings were meticulously transferred to a 3D-printed dual-ring biobot skeleton using a tweezer. Media was then changed to the differentiation media (DM^++^) consisting of ~98% DMEM supplemented with 2% ACA and 0.005% of insulin-like growth factor-I from mouse (IGF-1, Sigma Aldrich). 3D skeletal muscle rings were differentiated from myoblasts to myotubes and kept in DM^++^ until they mature. Media was changed daily, and the dual-ring biobots with muscle rings attached were kept in incubators at 37 °C and 5% CO_2_.

### B. Confocal imaging of muscle rings

Muscle rings were removed from the biobot structures at the end of each experiment and assessed with immunohistochemical staining and imaging. Tissue samples were rinsed with PBS and fixed in 4% v v^−1^ of paraformaldehyde for 20 min. To permeabilize the tissue samples, they were washed three times in PBS for 5 min and then incubated with 0.25% v v^−1^ Triton-X diluted PBS for 15 min. The muscle rings were then blocked and stored in 1% w v^−1^ bovine serum albumin (Sigma-Aldrich) at 4 °C overnight. The primary antibodies, mouse anti myosin heavy chain (MF-20) and rabbit anti *α*-actinin, were used to stain for myosin heavy chain and the sarcomere respectively with a 1:500 and 1:200 dilution ratio. The muscle samples were incubated overnight at 4 °C and then washed three times for 5 min before staining with secondary antibodies on the next day. The secondary antibodies, AlexaFluor-488 anti-rabbit and AlexaFour-568 anti-mouse (ThermoFisher), were used to stain *α*-actinin and MF-20 antibodies respectively. The samples were incubated overnight with DAPI to stain nuclei at 4°C. After washing with PBS three times, the LSM 700 was used for the confocal fluorescent imaging.

### C. Electrical stimulation of dual-ring biobots

In order to induce the contraction of 3D skeletal muscle rings, a function generator was used to depolarize the muscle tissue. The function generator ran at 20 V, and data collection was done over a series of frequencies including 1 Hz, 2 Hz, 3 Hz and 4 Hz. The dual-ring biobots were placed in a 35 mm dish with 4 mL of DMEM warmed to 37 °C in a water bath. The lid of the dish was fitted with 2 platinum (Pt) wires (20 mm) that ran lengthwise vertically across the dish (Figure S1a). Two Pt electrodes were connected to the function generator, creating a symmetric electric field perpendicular to the bots resulting in straight locomotion. A second lid was fabricated with two shorter Pt electrode (3 mm) to generate an asymmetric electric field for turning (Figure S1a). Prior to stimulation, the dish lids fabricated with Pt wires were submerged in 70% isopropanol for 2 min and submerged in PBS for 2 min for sterilization. Data was recorded through a stereomicroscope (MZ FL III, Leica Microsystems) with a field of view large enough to capture the entire dual-ring biobot. Video was captured at a frame rate of 10 f s^−1^ with a digital microscope camera (Flex, SPOT Imaging Solutions) and processed in ImageJ software (NIH).

### D. COMSOL^®^ modeling of electric field stimulation

To quantify the electric field strength that was applied to the dual-ring biobot, a 3D simulation model of the symmetric/asymmetric stimulation setup was designed using AC/DC electrostatic module (COMSOL^®^ Multiphysics 5.3a) (Figure S4). The domain and boundary conditions of each material were obtained from the literature[37, 38], and the electric conductivity of cell media (1.4 S m^−1^) was experimentally measured. To achieve satisfactory solving resolution, the maximum mesh element size (~9 · 10^−4^ mm) was set, which is smaller than the minimum displacement of the walker (~10^−3^ mm).

### E. Video tracking and force calculation

Force data was collected over the range of listed frequencies,1 Hz, 2 Hz, 3 Hz and 4 Hz via video capture with the bot pillars facing the camera. Locomotion of the bots was recorded with the pillars facing the bottom of the dish at each of the frequencies. Passive tension was collected through a still image captured with the bot laying on its side, and the bending of center beam was measured. The DMEM solution was changed in between each successive dual-ring biobot. The pillars’ deflection and locomotion were tracked using Tracker software.

Muscle force was calculated by the Euler-Bernoulli beam theory using angle of deflection with small angle approximation.

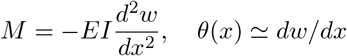

Angle of deflection was attained by 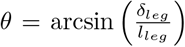, where *δ_leg_* is the deflected displacement per leg in side view and *l_leg_* the average of leg lengths due to the asymmetric structure of testbed. Likewise, the position of bending was selected as half of the beam length, x = L/2 and the moment caused by muscle force was written *M* = *F* × *l*, where *l* is the distance from beam to muscle location. As the result, muscle force was calculated in the form of

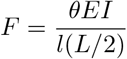

where I is the second moment of inertia of the beam and E stands for Young’s modulus, which was measured to be 350 kPa.

### F. Computational modeling and simulation

We computationally model the dual-ring biobot utilizing *Elastica*, an open-source software that we have developed for simulating complex, heterogeneous architectures made of soft slender bodies [30, 31]. These are modeled as Cosserat rods, a mathematical description that captures the dynamics in 3D space of 1D slender bodies, accounting for all modes of deformation (bending, twisting, stretching and shearing). A bio-hybrid walker is then modeled as an assembly of Cosserat rods specialized to muscular and scaffold elements. This rods assembly is constructed via appropriate boundary conditions to account for structural connectivity and dynamic interactions among rods. Furthermore, environmental effects such as gravity, buoyancy, surface friction and hydrodynamics are also characterized and implemented in simulation. Further details relative to our computational approach can be found in Supplementary Materials and in [30, 31]. The numerical solver *Elastica* is publicly available at https://www.cosseratrods.org.

## V. ACKNOWLEDGEMENTS

This study is jointly funded by NSF EFRI C3 SoRo #1830881 (M.G., R.B.), NSF CAREER #1846752 (M.G.), and Strategic Research Initiatives (SRI) program of the University of Illinois at Urbana-Champaign (M.G., R.B.). We also thank the Blue Waters project (OCI-0725070, ACI-1238993), a joint effort of the University of Illinois at Urbana-Champaign and its National Center for Supercomputing Applications, and the Extreme Science and Engineering Discovery Environment (XSEDE) Stampede2, supported by National Science Foundation grant number ACI-1548562, at the Texas Advanced Computing Center (TACC) through allocation TG-MCB190004. The authors thank Onur Aydin and Yukun Gong for reviewing various sections of the manuscript. We also thank Hunter Smith, and Elizabeth Pierson for helping with the CAD design and fabrication.

## Notes

### Competing Interest Statement

The authors have declared no competing interest.

